# Ultrahigh-Throughput Förster Resonance Energy Transfer-Activated Droplet Sorting for Terminator Polymerase Engineering

**DOI:** 10.64898/2025.12.18.695305

**Authors:** Hui Zhang, Qingqing Xie, Haoran Xu, Wei Li, Chuang Li, Liang Shen, Peiyi Wang, Xiangwen Yan, Xiang Li, Shiyue Zhou, Shitian Zhuo, Chuanyu Liu, Bo Teng, Yuliang Dong, Mengzhe Shen, Yue Zheng, Xun Xu

**Affiliations:** College of Life Sciences, University of Chinese Academy of Sciences, Beijing 100049, China; State Key Laboratory of Genome and Multi-omics Technologies, BGI Research, Shenzhen, 518083, China; State Key Laboratory of Genome and Multi-omics Technologies, BGI Research, Hangzhou 310030, China; BGI Research, Changzhou 213299, China

**Keywords:** Droplet-Based Microfluidic Screening, Protein Engineering, DNA polymerase, next-generation sequencing, modified nucleotide

## Abstract

Droplet microfluidic screening systems enable high-throughput, labor-saving enzyme directed evolution by employing fluorescence, absorbance, and Raman-activated sorting strategies for library screening. Förster resonance energy transfer (FRET) – a nanoscale technique for monitoring intramolecular/intermolecular conformational changes – is yet to be integrated into this process. We upgraded a single-channel sorter to a dual-channel one without redesigning the microscopy setup, which can monitor FRET signals during enzyme reactions in droplets at kilohertz rates. We applied this upgraded sorter to improve the incorporation efficiency of KOD DNA polymerase towards reversible terminator deoxyribonucleotides, a property crucial for its application in next-generation sequencing (NGS). Our data show that a single-round sorting can achieve 30-fold enrichment of active variants. Five KOD variants enabling 100-cycle single-end runs of DNA sequencing were identified using a novel cyclic reversible termination (CRT) substrate featuring a terminator group ≈ 5-fold bulkier than the classical azidomethyl moiety. We also engineered a metagenome-derived novel polymerase, and a variant achieving 90% terminator incorporation efficiency within 2 minutes was identified after two rounds of enrichment. In sum, we provide a practical setup for dual-channel FRET-based droplet sorting, and demonstrate its ability in terminator polymerases engineering, thereby broaden the scope of microfluidic applications.

## Introduction

Biological catalysts demonstrate notable advantages over traditional chemical catalysts in reaction efficiency, selectivity, and environmental compatibility, and are driving the transition of industrial processes towards sustainability. However, natural enzymes often exhibit limitations in substrate specificity and catalytic activity that cannot meet the needs of industry. Directed evolution accelerates natural evolutionary process through iterative cycles of gene diversification and variants library screening, enabling rapid engineering of biocatalysts for enhanced or entirely new functions in laboratory^[1]^. Yet screening assays remain a significant bottleneck due to substantial time requirements. Microfluidics-based screening addresses this by compartmentalizing reactions in water-in-oil emulsion droplets and screening in an ultrahigh-throughput way ^[2,3]^. In addition, this miniaturization maintains phenotype-genotype linkage, enabling reliable variants identification.

Diverse detection principles including fluorescence intensity^[2-4]^, absorbance^[5-7]^, Raman spectroscopy^[8,9]^, electrochemistry^[10,11]^ detection provide droplets-based directed evolution method a broader scope of applications. Currently, reported cases amenable to microfluidic-based enzyme screening include hydrolases^[12,13]^, oxidoreductases^[14,15]^, transferases^[16,17]^, lyases^[5,18]^, isomerases^[19,20]^, ligases^[20]^, translocases^[21]^, etc.. Fluorescence-based methods are highly favorable due to their high sensitivity (sub-nanomolar detection limits) and easy integration with microfluidics. However, the commonly applied fluorescence-based droplet screening methods remain challenging in either substrate design^[22-27]^ or signal-to-noise ratio (SNR) ^[5,28]^. To address the limitations and broaden the applicability of ultrahigh-throughput enzyme screening technologies, integrating novel detection schemes with microfluidic technology is essential.

FRET exhibits comparable throughput with fluorescence-based method^[29]^ and is a viable detection scheme candidate. FRET is a nonradiative distance-dependent (1–10 nm) energy transfer process where an excited donor fluorophore transfers energy to a proximal acceptor, enabling real-time monitoring of biomolecular interactions at nanoscale. Key applications of FRET span high-throughput discovery of small-molecule^[30,31]^, evaluation of anticancer compounds^[32]^, identification of heart-failure drug candidates in living cells^[33]^ and ultrahigh-throughput screening of antibody-secreting cells^[34,35]^. Despite FRET’s significant advantages, including low detection limits, microfluidic compatibility and intrinsic sensitivity to molecular distance /orientation changes, no FRET-activated droplet sorting system has been implemented for ultrahigh-throughput directed enzyme evolution to date.

Here in this work, we upgraded a single-channel fluorescence activated droplet sorter to dual-emission FRET based style by the combination of optical fiber embedded microfluidic chip. This design eliminates the need for external optical alignment and avoids microscopy setups redesign. In addition, we employed microcontroller unit (MCU) equipped with Kalman filtering algorithm^[36]^ to process the analog signals generated from photomultiplier tubes (PMTs) and to enhance SNR. Compared with field-programmable gate array (FPGA), MCU offers significantly lower costs, simplified development workflows, and superior power efficiency^[37]^. This setup enables FRET-based droplet sorting at rates of exceeding 1,000 droplets per second, which is 2-3 fold superior than reported FRET-based droplet sorting cases^[34,35]^.

As a model system for FRET-based enzyme screening, we selected the fluorescent reversible terminator polymerase (terminator polymerase), an essential tool enzyme for CRT-based NGS technology^[38]^. Unlike natural DNA polymerases performing continuous DNA synthesis, terminator polymerase exclusively incorporates reversible terminator nucleotides in a stepwise manner for single-base discrimination in DNA sequencing. Consequently, existing ultrahigh-throughput screening methods optimized for processive enzymes^[39-42]^ fail to accommodate its unique catalytic mechanism,while the conventional gel electrophoresis^[43]^ and microtiter plate assays^[44]^ restrict screening throughput to merely 10^2^-10^3^ variants per round due to serial processing limitations. Employing our in-house dual-channel FRET droplet sorting system, we firstly examined the terminator incorporation catalyzed by diverse DNA polymerase variants. A 30-fold enrichment of positive variant was achieved in a single sorting round with the sorting throughput of 10^3^ droplets per second (> 10^6^ mutants per day), 1,000 times faster than microtiter plate-based method. After two rounds, multiple KOD polymerase variants capable of incorporating the sterically challenging sequencing substrate 3’-O-2-[1-(ethyldisulfanyl) ethyl] benzene-1-carbaldehyde nucleotide (namely SSEB-dNTPs), which has a 5-fold bulkier blocking group than the classical 3’-O-azidomethyl reversible terminator, were obtained. Five KOD variants were functionally validated on the BGISEQ-500 sequencing platform using SSEB-dNTPs as CRT substrate, achieving Q30 score > 80%. Then, to rapidly engineer a newly discovered polymerase (∼ 40% sequence identity to the known family B polymerase) for NGS application, we screened a 16,000-member mutagenesis library via ultrahigh-throughput sorting. A variant exhibiting 90% substrate-to-product conversion within 2 minutes was identified after only two sorting rounds. Amino acid frequency analysis revealed significant enrichment of A/G/S/V/T substitutions at Y516-a residue homologous to Y409 in KOD polymerase and Y412 in Vent polymerase. These smaller side-chain substitutions confirm that Y516 in the new polymerase acts as a steric gate for nucleotide sugar recognition^[45]^.

In summary, our work establishes a robust dual-channel FRET droplet sorting platform and demonstrates its efficacy in engineering superior-performance terminator polymerases for NGS, thereby expanding microfluidics-driven directed evolution in biocatalysts development.

## Results and Discussion

### Design of a FRET-based Microdroplets Screening Platform

To achieve ultrahigh-throughput screening of microdroplets based on FRET signal at single cell level, a sensitive droplet-based optofluidic system was combined with a MCU-based signal processing program for the detection and sorting of donor–acceptor fluorophore pair (Figure 1A). To enable easy and cost-effective FRET signal acquisition within microdroplets, we integrated an optical fiber probe (50-μm core diameter, 0.22 NA UV/VIS patch cord) directly into the microfluidic chip. The optical fiber is 60 μm away from the flow channel. This design eliminates external optical alignment complexity, bypassing the need for microscopy setups redesign. By leveraging optical fiber transmission, background noise from excitation light is minimized. Furthermore, a PDMS injection outlet was integrated into the fiber-guide channel to exhaust air between the fiber and PDMS wall, eliminating the change in refractive indices and reducing light loss. A 532 nm laser beam was projected onto the microfluidic device via a 20× objective. As droplets traversed the excitation beam, emitted donor fluorescence was collected by the same objective and transmitted through a series of dichroic mirrors to a photomultiplier tube (PMT) equipped with a 585/29 nm bandpass filter. Acceptor fluorescence was collected by an optic fiber embedded within the chip and transmitted to a second PMT with a 681/24 nm bandpass filter.

**Figure 1.**
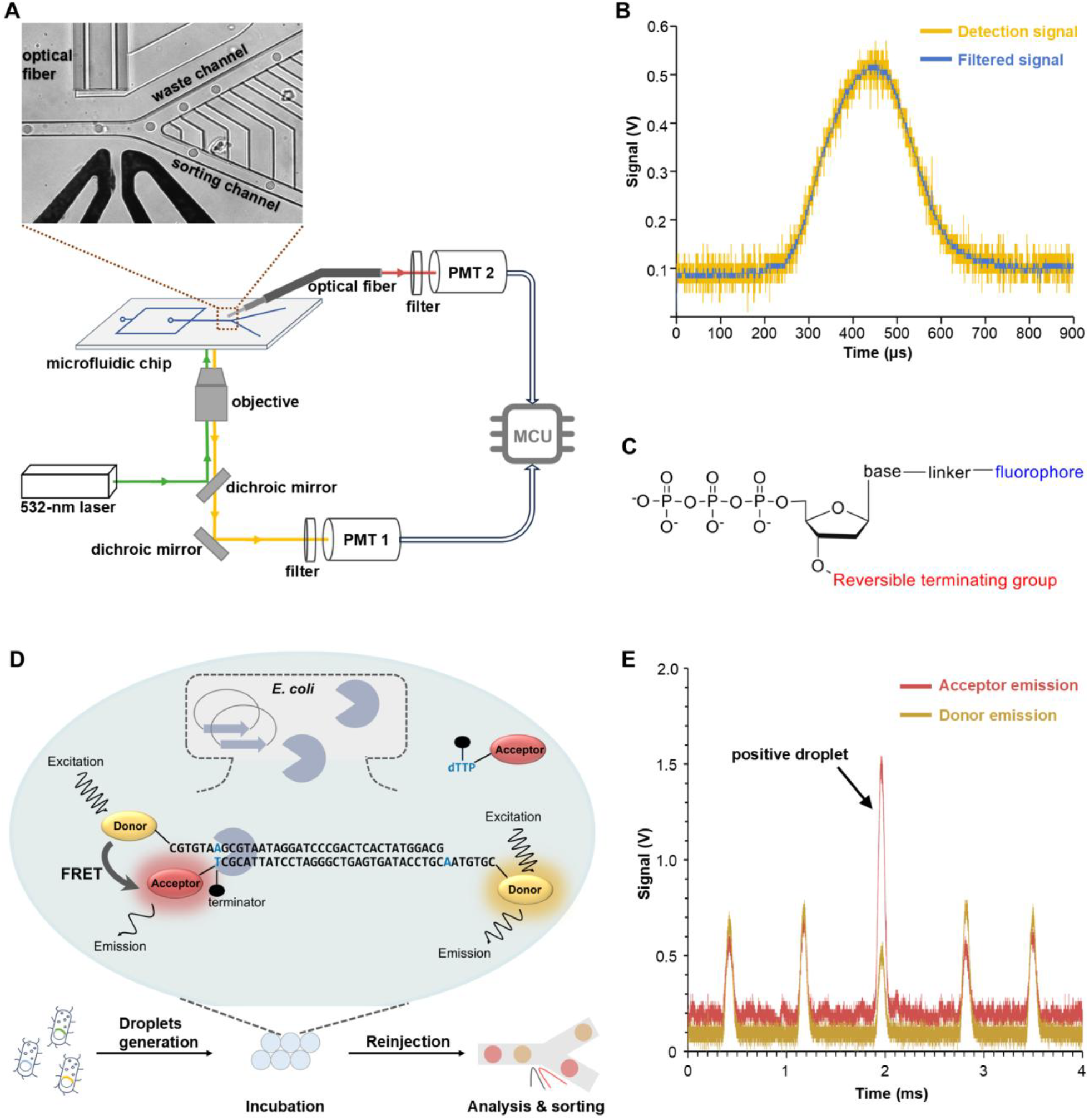
Optical fiber-based fluorescence detection combined with MCU-based signal processing enable robust analysis of FRET signals within microdroplets. A) Top: Image of droplets flowing through the sorting region, showing optical fiber, electrodes and sorting channel. Bottom: Schematic of the optical setup featuring optical fiber and objective for fluorescence detection and MCU for signal processing. B) Detection signal trace(yellow) and the filtered signal trace (blue) obtained by applying Kalman filter. C) Structure schematic of a reversible terminator used in NGS. The reversible terminator features a fluorophore (blue) linked to the nucleobase via a cleavable linker and a reversible terminating group at the 3’-OH position. D) Workflow of FRET-based microdroplets screening and enzymatic activity assay for terminator-incorporating polymerase. Polymerase expressing cells are encapsulated in 5 pL droplets with reversible terminator activity assay reagents. After incubation, droplets containing polymerase that can add acceptor labeled reversible terminator to donor labeled DNA primer-template complex will be sorted through the FRET-induced acceptor fluorescence signal increase and donor fluorescence signal decrease. E) Time series recording of an artificially mixed droplet library during a 4 ms sorting window, showing 5 droplets signals. The third droplet showing increased acceptor emission signal and decreased donor emission signal is a positve droplet. Yellow: FRET-correlated donor fluorescence signals; Red: acceptor fluorescence signals.

PMT generates thermal noise and shot noise during operation, while external environmental interference introduces vibration noise, and circuit noise is inherent in the analog-to-digital converter (ADC) sampling process for optical signal sensing. To enhance the SNR and relax stringent environmental requirements, we implemented a Kalman filtering algorithm in C language on MCU controller. This algorithm utilizes the “prediction” step to estimate the next state through system modeling (even without new measurements), followed by the “update” step that fuses this prediction with actual ADC readings, resulting in high-fidelity state estimation outcomes (Figure 1B). This dual-stage mechanism significantly improves the system’s responsiveness to signal transients and markedly reduces latency effects, with its operational principle illustrated below.

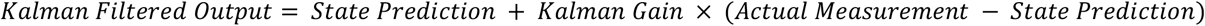

To demonstrate the efficacy of our FRET-based microdroplets screening platform for practical applications, the enzymatic activity of terminator polymerase was quantified within individual droplets. The terminator employed in CRT-based NGS workflows contains a 3’-OH blocking group (such as 3’-O-azidomethyl) and a base-linked fluorophore (Figure 1C), which impose steric hindrance to the active site of wild-type DNA polymerases, necessitating the use of engineered DNA polymerases for efficient incorporation. A previously described microplate-based terminator incorporation efficiency assay method^[44]^ was applied in our microdroplet system (Figure 1D). In brief, acceptor and donor fluorophores were conjugated to the base of nucleotide and the 5’ termini of DNA template, respectively. By the catalyzing of an active DNA polymerase, a terminator nucleotide is incorporated into the 3’ termini of the DNA primer, bringing two fluorophores within Förster resonance energy transfer (FRET) range (<10 nm). The FRET is triggered and the resulting change in Cy5 emission signal serves as polymerase activity indicator. In contrast to microplate-based assays employing purified enzyme and time-resolved acceptor fluorescence signal detection, our droplet-based approach directly compartmentalizes enzyme-expressing cell with fluorogenic reagents into water-in-oil microdroplets (20 μm in diameter) and employs endpoint signal detection to achieve ultrahigh-throughput enzyme screening. A potential limitation of this approach is that endogenous host DNA polymerases and exonucleases might catalyze side reactions, which could interfere with the FRET signals specifically generated by the target enzyme’s activity. To eliminate the side reactions, we optimized the droplet incubation procedure. After emulsification, the droplets were first incubated at 90 °C for 5 minutes to inactivate endogenous enzymes and lyse the cells, and then incubated at 58 °C for the terminator incorporation reaction. The limited availability of polymerase within single cell can lead to FRET signals being drowned out by background fluorescence. This arises due to the partial excitation of the Cy5 acceptor fluorophore from the unreacted substrate molecules by the 532 nm laser and the collection of emission from the directly excited Cy3 donor fluorophore within the 681/24 nm filter bandwidth. To mitigate the background noise resulting from the direct excitation, we minimized the substrate concentration to the lowest level compatible with the detection sensitivity of our platform. Furthermore, to enhance the reliability of signal detection and maximize the positive rate, we simultaneously monitored the reduction in donor fluorescence intensity and the corresponding increase in acceptor fluorescence intensity, ensuring robust identification of polymerization events.

A KOD polymerase variant Mut_C2 developed by our laboratory^[44]^ was used as demo to validate our FRET-based microdroplets screening platform. The Mut_C2 expressing *E. coli* cells were co-encapsulated with Cy5-labeled 3’-O-azidomethyl-dTTP and Cy3-labeled primer-template complex. The incubated droplets were subjected to FRET-based droplets analysis at kilohertz rates, real-time signals from droplets were recorded with oscilloscope (Figure 1E). Droplets exhibiting simultaneous donor fluorescence signal decay and acceptor fluorescence signal enhancement were classified as hits.

### Validation of The Platform for Terminator Polymerases Screening

We next evaluated the detection sensitivity of our FRET-based microdroplets screening platform by quantifying the correlation between the enzymatic activity of active polymerase and the FRET signals within individual droplets. We constructed a demo KOD polymerase mutant (KOD-M) contained two mutations (Y409A, S451T) from our previous engineering efforts^[44,25]^ and three mutations (D141A, E143A, A485L) from Therminator DNA polymerase^[46]^. Figure 2A displays scatterplots of FRET fluorescence signals obtained from three distinct droplet populations containing different concentrations of purified KOD_M polymerase (0 μM, 0.01 μM, and 0.2 μM, respectively) and identical concentration of Cy5-labeled 3’-O-azidomethyl-dTTP and Cy3-labeled primer-template complex in 1× Thermopol buffer (NEB, 20 mM Tris–HCl pH 8.8 at 25°C, 10 mM (NH4)_2_SO_4_, 10 mM KCl, 2 mM MgSO_4_, 0.1% Triton X-100). As incubated droplets sequentially traverse the detection region of the microfluidic chip, real-time optical signals are continuously acquired. Droplets containing active polymerase exhibit an obvious increase in acceptor (Cy5) fluorescence intensity accompanied by a corresponding decrease in donor (Cy3) fluorescence intensity. The fluorescence signal profiles of enzyme-active droplets are significantly different from the empty droplets, enabling clear discrimination during ultrahigh-throughput screening. This result demonstrate that our detection system exhibits sufficient sensitivity to resolve polymerization signals generated by KOD_M polymerase at concentrations as low as 0.01 μM (equivalent to 2500 enzyme molecules per droplet). This corresponds to a limit of detection (LOD) comparable to the level of heterologous protein expression in a single cell. This finding suggested that using the droplet-based FRET signal to assess the polymerization activity of variant towards terminator is reliable.

**Figure 2.**
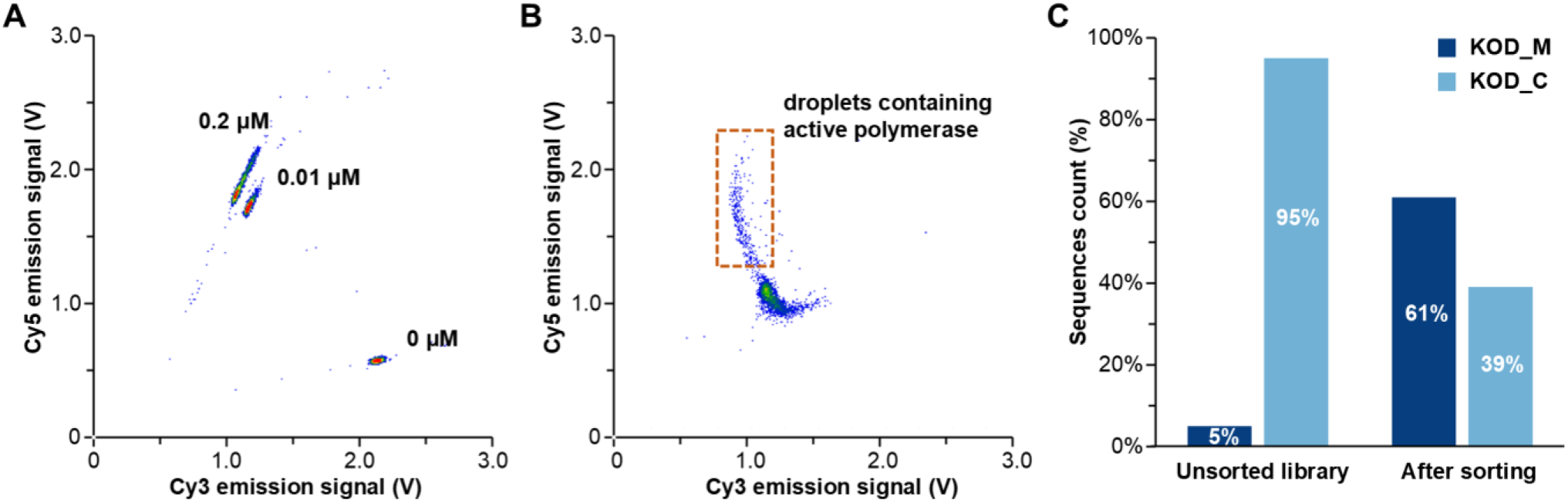
Performance evaluation of the FRET-based microdroplets screening platform. A) Scatterplots of FRET signal from droplets containing Cy5-labeled 3’-O-azidomethyl-dTTP and Cy3-labeled primer-template complex supplemented with varying concentrations of KOD_M polymerase (0 μM, 0.01 μM, and 0.2 μM). B) Scatterplots of FRET signal of droplets containing a mixed library of KOD_M variant (active) and KOD_C variant (inactive) with Cy5-labeled 3’-O-azidomethyl-dTTP and Cy3-labeled primer-template complex in 1x Thermopol buffer. The boxed region indicates the subpopulation enriched. C) KOD_M cells were enriched 30 folds after sorting, confirming successful enrichment of active polymerase variants from the mixed library.

To evaluate the sorting efficiency of our system for discriminating between *E. coli* cells expressing active and inactive polymerase variants at single-cell level, we co-encapsulated strains harboring either an inactive KOD_C polymerase (KOD exonuclease deficient mutant D141A/E143A) or an active KOD_M polymerase along with identical substrates into microdroplets (λ=0.1) suspended in 1× Thermopol buffer. Following incubation, the droplets were analyzed on our screening platform to record both the Cy3 and the Cy5 signals. The droplets within the defined frame were sorted out (Figure 2B). Both unsorted and sorted droplets were collected, from which the polymerase variants’ genes were cloned for Sanger sequencing to evaluate the sorting efficiency (100 clones for each group). Our FRET-based microdroplets screening platform successfully enriched active KOD_M polymerase from a mixed library comprising predominantly inactive variants, achieving a 30-fold enrichment following a single round of sorting (Figure 2B). The enrichment factor was determined by applying the following formula: (KOD_M sorted/KOD_C sorted) /(KOD_M unsorted/KOD_C unsorted). These results demonstrate that our platform possesses the requisite sensitivity and accuracy to screen for polymerases capable of efficient terminator incorporation.

### Directed Evolution of Terminator Polymerase for Novel Sequencing Chemistry

The efficient primer-dependent incorporation of fluorescent reversible terminator deoxyribonucleotides by engineered DNA polymerases presents a fundamental challenge for the development of CRT-based NGS technology. Our FRET-based microdroplets screening platform was applied in the ultrahigh-throughput KOD DNA polymerase engineering, aiming to alter its substrate specificity for efficient incorporation of a novel reported terminator^[47]^. This terminator, abbreviated as SSEB-dNTPs, features a 2-[1-(ethyldisulfanyl) ethyl] benzene-1-carbaldehyde group capping the 3’-OH. Compared to 3’-O-azidomethyl-dNTPs, SSEB-dNTPs harbors a 5-fold bulkier 3’-O-blocking group (Figure 3A) which imposes significantly greater steric hindrance within the active site of DNA polymerases. This enhanced steric hindrance restricts the polymerization activity of the existing KOD_M polymerase, leading to inefficient incorporation of SSEB-dNTPs in NGS. Residues L408/Y409/P410 located in close proximity to the 3’-OH group^[48]^ (Figure 3B) were simultaneously mutated. By employing VNK-degenerate primers to exclude termination codon mutants, the proportion of functional clones is significantly increased, leading to a more efficient library screening process. In the primary screening round, the original library (comprising 4,096 variants) was co-encapsulated into droplets with the polymerase activity assay mixture and subjected to droplet-based optical polymerase sorting (DrOPS) ^[42]^. This initial step effectively excludes mutants that either expressed insoluble protein or lacked fundamental polymerase activity. Subsequently, the enriched mutant library from the first round was retrieved, co-encapsulated with Cy5-labeled SSEB-dTTP and Cy3-labeled primer-template complex in 1x Thermopol buffer, and subjected to a more stringent FRET-based droplets sorting. To ensure comprehensive coverage, the entire mutant library was sampled by screening 2.54 million droplets (>155-fold oversampling at 25% droplet occupancy), and 59,000 droplets were selected for further analysis, corresponding to the top 2.3% of the brightest droplets. The genes expressing polymerase variants from sorted droplets were extracted, amplified and subcloned. To evaluate screening efficiency, 45 clones randomly picked from each of the original library, the primary sorting output, and the second sorting output were analyzed using a lysate-based microtiter plate assay. The results demonstrated successive enrichment of active clones and effective selection of high-performance variants throughout the sorting process (Figure 3C).

**Figure 3.**
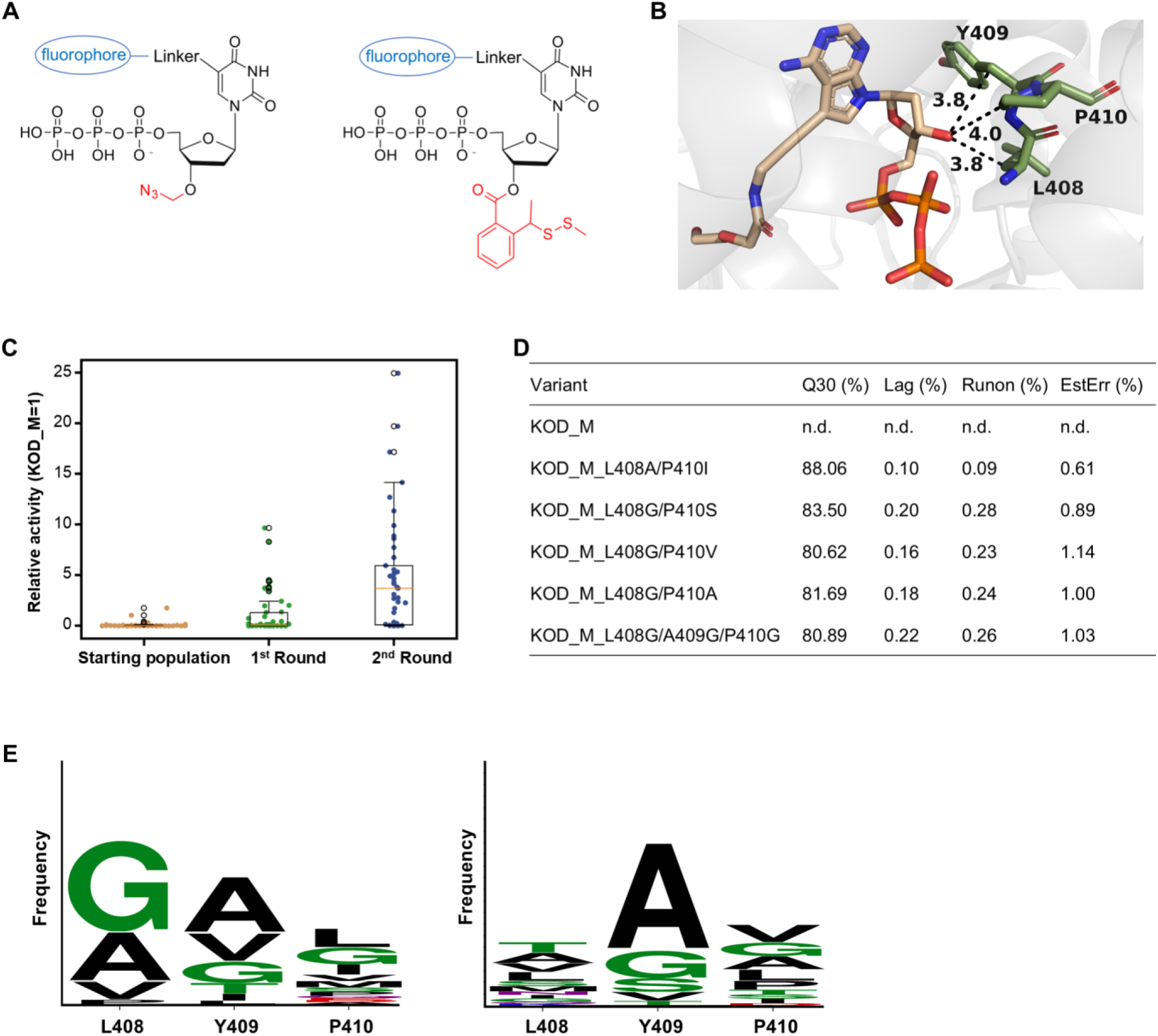
KOD DNA polymerase were engineered with tailored substrate preferences for massive parallel sequencing application. A) Structure schematic of fluorescently labeled 3’-O-azidomethyl-dTTP (Left) and SSEB-dTTP (Right). B) Crystal structure of KOD DNA polymerase in a closed ternary complex with 7-deaza-7-(2-(2-hydroxyethoxy)-N-(prop-2-yn-1-yl) acetamide)-2-dATP (PDB ID: 6Q4T). L408, Y409 and P410 are located in close proximity to the 3’-hydroxyl group of 7-deaza-modified adenosine in the crystal structure. C) A representative comparison of the reversible terminator polymerization activity distribution among the starting population, the population after the first round of sorting with natural dNTPs, and the population after the second round of sorting with Cy5-labeled SSEB-dTTP, displaying the enrichment of variants that have increased activities. 45 clones were randomly selected from each group for analysis. D) The sequencing metrics (Q30, Lag, Runon, EstErr) of KOD_M and selected variants were evaluated on the BGISEQ-500 sequencing platform under the condition of using fluorescently labeled SSEB-dNTPs for single-end 100-cycle. E) The residues proportions at 408/409/410 of 36 most active KOD mutants for SSEB-dNTPs incorporation (left) and 3’-O-azidomethyl-dNTPs incorporation (right)

The applicability of the sorted polymerase variants was evaluated within massive parallel sequencing workflows on the BGISEQ-500 sequencing platform^[49]^ utilizing fluorescently labeled SSEB-dNTPs as the CRT substrate. The top 30 variants exhibiting the highest activity in the microtiter plate assay were subjected to expression and purification. Sequencing quantifying metrics including Q30, lag, runon and estimate error rate (EstErr) were used to benchmark their performance in commercial sequencing instrument. KOD_M polymerase failed to complete the run, exhibiting a drastic reduction in DNA nanoball (DNB) intensity and premature termination due to severely impaired recognition and incorporation of the challenging substrate. In contrast, under identical sequencing conditions, five variants featuring smaller side-chain substitutions at positions 408 and 409 (Ala and Gly) exhibited enhanced sequencing durability, achieving 100-cycle single-end reads with a Q30 score exceeding 80%. Four of them exhibited lag values passing the quality control criteria of 0.20%, confirming minimal signal decay and high synchrony throughout the 100-cycle sequencing workflow. Among them, the mutant KOD_M_L408A/P410I performs the best, with a Q30 value reaching 88%, a lag value as low as 0.10%, a runon value as low as 0.09%, and an EstErr value of 0.61%.

For ultrahigh-throughput screening of terminator polymerase using microdroplets, only microliter volumes of sequencing substrates were consumed. Furthermore, because the microdroplets screening discarded the vast majority of mutants exhibiting no or low activity, only 10^2^ mutants required validation via the slower and more costly microplate-based assays and sequencer-based methods. Among these, mutants with a Q30 value greater than 80% were successfully identified. Overall, the integration of droplet microfluidics and FRET detection in our platform provides an effective solution for engineering high-performance polymerases tailored to NGS chemistries, with demonstrated success in reducing screening costs and accelerating enzyme engineering cycles from months to weeks.

Meanwhile, the amino acid proportions at 408/409/410 in the active mutants group were calculated to give a clear view on which amino acids confer activity and tend to be enriched during different terminator polymerase evolution. The above-mentioned KOD mutant library was subjected to FRET-based droplets sorting with Cy5-labeled 3’-O-azidomethyl-dTTP as terminator. Polymerase gene from sorted droplets was extracted, cloned and validated by microplate reader. 36 high-performance variants each from the SSEB and azidomethyl terminator polymerase group was sequenced to analyze the frequency of amino acids encountered at investigated position (Figure 3E). From the sequence logo, we concluded that Y409 is in general replaced with small amino acids as a result of steric hindrance caused by the 3’-OH modification to enhance the terminator incorporation efficiency. Additional small amino acid substitutions for L408 were necessary for incorporating SSEB terminator efficiently. While more alternative residues found at position 408 for azidomethyl terminator polymerase, indicating the SSEB substituent in the incoming nucleoside triphosphate would sterically clash with the residues of both 408 and 409. Exceptionally, variants with Y409I or Y409L substitution (KOD_M_L408A/A409I/P410L, KOD_M_L408G/A409I/P410G) incorporate SSEB-dNTPs efficiently which achieved 100-cycle single-end reads with a Q30 score 77% and 64%. This discovery conflicts with the previous studies that reducing the size of the side chain at the 409 (from Y to V/A/G mutation) will create polymerases that can incorporates modified nucleotides^[45,50,51]^, as the Tyr in the conserved LYP motif act as a “steric gate” for sugar discrimination for B family polymerase. The reason why Leu and Ile substitution of KOD Y409 result enhanced SSEB-dNTPs polymerization activity may be attributed to the effects of 408 and 410 residues that maintain the proper orientation of the nucleotide and the side chain of the steric gate residue thereby enabling SSEB-dNTPs recognition.

### Rapid engineering of novel DNA polymerases

Having demonstrated the capability of our FRET-based microdroplets screening platform to engineer known polymerase for novel substrate incorporation (as with KOD DNA polymerase), we next explored its applicability in accelerating the functional optimization of four newly mined DNA polymerases for the incorporation of 3’-O-azidomethyl-modified terminator. These four novel DNA polymerases (designated 48°N^[52]^, 27°N^[53]^, 4°S^[53]^, and 22°S^[54]^), belonging to the polB family, were mined from the deep-sea hydrothermal sediment metagenomic data^[55]^. They exhibit a melting temperature (Tm) exceeding 90 °C and possess the ability to amplify DNA fragments by PCR and perform error correction via 3’–5’ exonuclease activity, similar to the commercial DNA polymerases derived from Thermococcus kodakarensis KOD1, Thermococcus sp. 9°N-7, Thermococcus gorgonarius, and Desulfurococcus sp. Tok, Pyrococcus abyssi, and Pyrococcus GB-D. Notably, these four novel polB polymerases exhibit relatively low sequence identity (approximately 40%) to the aforementioned commercial polymerases (Figure 4A), suggesting that they are phylogenetically distinct. In addition, the protein structures of these four polymerases were predicted using AlphaFold2^[56]^ and aligned to the experimental structure of KOD DNA polyerase (PDB ID: 4k8z) via the Universal Structural alignment (US-align) webserver^[57]^. The resulting TM-score exceeding 0.9 indicates that, despite the low sequence identity (∼40%), the three-dimensional structures of these novel polymerases are highly conserved and functional homologous, thereby explaining their similar catalytic activities. In summary, despite their low sequence homology, the high structural similarity and deep-sea hydrothermal vent origin of these enzymes suggest significant potential for engineering them into terminator polymerase.

**Figure 4.**
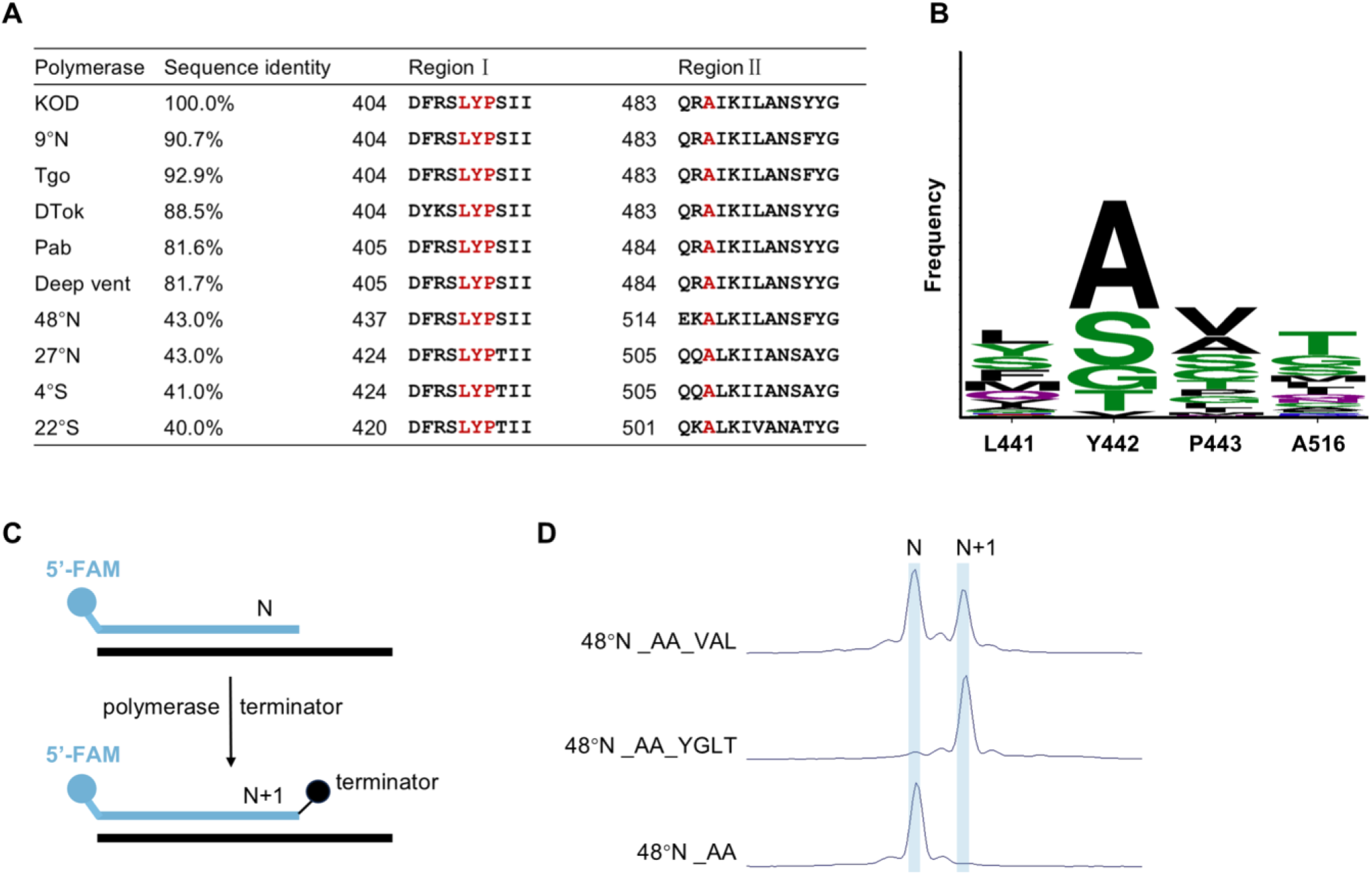
Engineering a novel family B polymerase with enhanced catalytic activity for terminator incorporation via a FRET-based microdroplets screening platform and combinatorial mutagenesis. A) Sequence alignment reveals low sequence identity between novel family B DNA polymerases and canonical family B polymerases. Four key residues (highlighted in red) within the polymerase domain were selected for combinatorial saturation mutagenesis. B) Frequency of amino acids encountered at the investigated positions for a total of 81 active clones sequenced after sorting with Cy5-labeled 3’-O-azidomethyl-dTTP. C) Schematic illustration of the terminator incorporation assay. D) 3’-O-azidomethyl-dTTP incorporation efficiency at 58 °C for 2 min of the top-performing 48°N variant, 48°N_AA_YGLT, as revolved by capillary gel electrophoresis. The primer (N) and the primer extended by one incorporated reversible terminator (N+1) are indicated. Incorporation results of 48°N_AA_VAL and exonuclease-deficient mutant 48°N_AA (48°N_D169A/E171A) under identical reaction conditions are showed for comparison.

To construct mutants for improving catalytic activity towards terminator, candidate mutation sites were first selected based on multiple sequence alignment of these four novel polymerases with typical family B DNA polymerases (Figure 4A). To prevent excision of newly incorporated terminator, 3’,5’-exonuclease-silencing mutations (equivalent to D141A and E143A in KOD_M) were introduced. KOD_M_L408V identified through droplet-based screening demonstrated superior activity towards 3’-O-azidomethyl-modified terminator — achieving a Q40 value reaching 94.22% and a Lag value of 0.09% in single-end 100-cycle sequencing. We introduced these beneficial mutations L408V/Y409A/A485L of KOD to the corresponding residues of 48°N, 27°N, 4°S, and 22°S. These resulting variants were subjected to activity assay with microplate reader-based method, and only the 48°N_D169A/E171A/L441V/Y442A/A516L (designated 48°N_AA_VAL) variant exhibited polymerization activity towards 3’-O-azidomethyl-modified terminators. We performed a fluorescent primer extension assay (Figure 4B) to quantify the nucleotide incorporation efficiency of 48°N_AA_VAL using fluorescence capillary gel electrophoresis method^[58]^. The reaction mixture, consisting of 100 nM purified polymerase and 30 nM FAM-labeled primer-template complex in 1x Thermopol buffer, was initiated by adding 1 μM 3’-O-azidomethyl-dTTP, followed by incubation, reactions were terminated with 100 mM EDTA. Capillary electrophoresis (CE) was conducted using an Applied Biosystems 3730xl Genetic Analyzer, with fluorescent signals quantified via GeneMarker software (SoftGenetics). CE analysis revealed that only 40% of the substrates were polymerized within 2 minutes under the reaction condition of 58 °C.

Furthermore, we combined focused, combinatorial libraries with the ultrahigh-throughput of droplet microfluidics to identify high-performance terminator polymerase. Four residues located within the conserved motifs A and B^[45]^ of the polymerase activity center were simultaneously subjected to saturation mutagenesis using NNK degenerate codon to encode all 20 canonical amino acids on the basis of 48°N, 27°N, 4°S, and 22°S exonuclease-deficient mutants. Four mutant libraries, each consisting of 16,000 (20^4^) variants, were individually expressed and screened using our FRET-based microdroplets screening platform in the presence of a 3’-O-azidomethyl-modified terminator substrate within 1x Thermopol reaction buffer. Among these, only the 48°N polymerase mutant library exhibited a droplet signal scatter plot comparable to that in Figure 2B. This demonstrates that the 48°N mutant library contains functional mutants, indicating the 48°N polymerase’s potential for engineering into a high-performance terminator polymerase for NGS technology. Then, the 48°N polymerase mutant library was subjected to DrOPS using standard nucleotides as substrates. The entire library was sampled by screening 8.29 million droplets (≈130-fold oversampling at 25% droplet occupancy), from which 78,000 droplets were selected. The enriched library was subsequently amplified, expressed, and underwent a more stringent round of FRET-based droplets sorting with 3’-O-azidomethyl-modified terminator as substrate. In this secondary screening, 3.08 million droplets were screened, and 104,000 droplets were selected. Following droplets sorting, a microtiter-plate-based polymerization activity assay was performed on 95 randomly selected clones. Active clones were recovered and subjected to Sanger sequencing. 85.3% (81/95) of the randomly selected clones showed activity towards 3’-O-azidomethyl-modified terminator. Sanger sequencing results of 81 active mutants revealed that substitution of Y442 with amino acids bearing smaller sidechain significantly impaired terminator incorporation activity, indicating the critical role of this residue in catalytic function (Figure 4B).

Using fluorescence capillary gel electrophoresis-based incorporation efficiency assay, we identified a novel mutant 48°N_D169A/E171A/L441Y/Y442G/P443L/A516T (designated 48°N_AA_YGLT), which achieved 90% substrate-to-product conversion within 2 minutes under the specified conditions (Figure 4D). For comparative purposes, incorporation results of both 48°N_AA_VAL and the exonuclease-deficient mutant 48°N_D169A/E171A (48°N_AA) were also presented (figure 4D).

Our novel polymerase engineering process demonstrates that rational site-directed mutagenesis of the novel polymerase based on the prior-knowledge of KOD, has limited effect on improving its activity. On the contrast, ultrahigh-throughput screening technology enables rapid assessment of the existence of postive variants within libraries comprising hundreds of thousands of variants, as well as precise identification of variants with significantly enhanced catalytic activity. Additionally, the engineered 48°N DNA polymerase demonstrates strong activity towards modified nucleotides, highlighting its potential as a tool for the synthesis of artificial genetic polymers (XNAs) ^[59]^.

## Conclusion

We present an integrated setup for ultrahigh-throughput FRET signal detection in microdroplets utilizing an embedded fiber-optic microfluidic chip combined with an MCU-implemented Kalman filtering algorithm. This design simplifies the optical architecture and significantly enhances the SNR with a screening throughput of over 1,000 droplets per second. In our research, compartmentalized enzymatic reactions leading to changes in FRET signal was systematically characterized, broadening the range of assayable catalytic reactions within droplet. This FRET-based microdroplets screening platform was then combined with structure-guided design to reprogram the substrate specificity of KOD DNA polymerase. This approach yielded several variants that were successfully deployed in a commercial sequencing platform. Furthermore, a newly mined polymerase 48°N which shares low sequence identity with known family B DNA polymerases was rapidly engineered for efficient terminator incorporation. Moreover, a 48°N mutant achieving 90% substrate-to-product conversion within 2 minutes under the specified reaction conditions was obtained. Owing to the ultrahigh-throughput and accuracy of our screening platform, amino acid frequency calculation at investigated positions among active variants can be quantified, revealing residue-specific contributions to activity. Our analysis revealed that tyrosine at motif A of polB is frequently substituted by smaller amino acids to alleviate steric hindrance caused by the 3’-OH modification. Moreover, additional substitutions of leucine with small residues at motif A proved necessary for efficient accommodation of substrates with bulkier 3’-OH blocking group.

Inspired by these findings, we propose that mining and engineering of polymerases with low sequence identity to known counterparts is a promising strategy for discovering enzyme candidates, thereby paving the way for innovations in sequencing technology. For the development of scarless CRT-based NGS, the presence of bulky fluorescent protecting groups at the 3’-OH position remains a major barrier to fast and efficient incorporation of dNTPs^[60]^. By utilizing our ultrahigh-throughput terminator polymerase screening method, it will be promising to identify proper polymerases capable of efficiently incorporating nucleotides with 3’-O-blocking groups that are both photolabile and fluorescently active^[61]^,promoting the development of next-generation sequencing chemistry.

## Acknowledgements

This work was supported by National Key Research and Development Program of China (2023YFC3402901).

