## Supplementary material for "Ultrahigh-Throughput Förster Resonance Energy Transfer-Activated Droplet Sorting for Terminator Polymerase Engineering": Suplementary method & figure & table

### *Supplementary Information*

#### Materials and Methods

#### Materials

All strains and vectors utilized in this work are summarized in Table S1. The oligonucleotides were devised and created by GCATbio (Changzhou, China) (Table S2).

#### Microfluidics device design and fabrication

Microfluidic chips were manufactured by polydimethylsiloxane (PDMS, Dow Corning) using SU8-on-Silicon-wafer masters (Hicomp Microtech). The droplet sorting chip is a two-layered microfluidics chip. The first layer is designed for droplets reinjection with a height of 20 μm, and the second layer contains sorting electrodes filled with fusible Indium-lead-tin alloy (Wo Chang metal), embedded optic fiber (Edmund Optics) with a height of 110 μm and part of channel. The design of the droplet sorting chip is shown in Supplementary Figure S1b.

#### Site-directed mutagenesis of polymerase variants used in this study

PD441 expression vector harboring the wild-type KOD gene and pET28a expression vector harboring the wild-type 48°N gene are lab stock. Mutations were introduced by PCR using high-fidelity Pfu DNA polymerase (Promega) and mismatch containing primers (Table S4). PCR product was digested with DpnI (New England Biolabs) to remove methylated and hemimethylated DNA from the parent template before transformation of E. coli DH5a. Single colonies were picked for plasmid isolation and sanger sequencing.

#### Generation of mutagenic plasmid libraries used in this study

Gene for mutant libraries construction were generated and multiplied using gene specific primers containing degenerate codon. The PCR consisted of 0.3 μM each of primers (Table S2), 1x KAPA HiFi HotStart ReadyMix (Roche), 5 ng template, in a 100 μL reaction. The reaction and droplet generation oil (QX200™ Droplet Generation Oil for EvaGreen, Biorad) were mixed and vortexed to generate droplet for emulsion PCR which avoiding bias. The droplets generated by vortexing were aliquoted into several PCR tubes. Light mineral oil is added to PCR tubes to prevent droplet coalescence caused by evaporation during thermal cycling. Following PCR amplification, transfer as many droplets as possible into a new Eppendorf tube. Then, add an equal volume of 1H,1H,2H,2H - perfluorooctanol (Sigma - Aldrich) to the tube. Vortex the tube for approximately 30 s and centrifuge it at 2000 r.c.f. for 1 min. The top aqueous layer containing the variant gene was purified on an agarose gel. Vector was amplified using high-fidelity Pfu DNA polymerase (Promega) and primers with overhangs complementary to gene. The PCR product was digested with DpnI and verified for correct size on the agarose gel, cut from the gel and purified by DNA gel purification kit (Tiangen). 3:1 molar ratio purified gene/vector was mixed with equal volume of Gibson Assembly Master Mix (New England Biolabs), following 15 minutes incubation at 50 °C. Mixture was purified by DNA Clean & Concentrator-5 (Zymo Research) and transferred into E. coli DH5a electrocompetent cells. The colonies were scraped with a sterile L-shaped spreader after pipetting LB medium onto the overnight plate. Mutagenic plasmid DNA was extracted from the collected cells by plasmid miniprep kit (Tiangen).

#### Preparation of bacterial suspension for droplets generation

Mutagenic plasmid was transferred into E. coli BL21(DE3) electrocompetent cells by electroporation, following standard procedures. A serial dilution of transformation suspension was spread onto LB/agar plates and incubated to calculate the total number of transformants. For each transformation, 20 clones were randomly picked and sequenced to evaluate the diversity of constructed library (Figure S3). Fresh transformations were recommended to maximize the amount and quality of recombinant polymerase expressed which is critical to identifying new variants with improved activity[62]. After 12 hours growth, the colonies were washed out and suspended with LB medium. LB-kanamycin (25mL, 50μg/mL) was inoculated with freshly transformed BL21(DE3) cells to a starting OD600 = 0.05 and incubated for ≈ 2 h at 37 °C and 280 rpm (until an OD600 = 0.6 was reached). KOD and 48°N expression were induced by the addition of IPTG (0.5 mM final concentration) and expression for 12 hours at 25 °C and 280 rpm. After expression, 1 mL of cell culture was centrifuged for 5min at 2,000 r.c.f. and the supernatant discarded. The cells were washed three times with 1× ThermoPol buffer (New England Biolabs). The rinsed bacterial pellet was re-suspended in 1mL 1× ThermoPol buffer and the absorbance was measured at 600 nm.

**Cell compartmentalization in droplets for activity selection**

Cells were diluted to enable encapsulation at occupancies of 0.1 cells per droplet according to the assumption that 1ml of E. coli suspension at an OD600 value of 1.0 contains $5\times10^8$ cells. Using a microfluidic device (Figure S1a), the cells and equal volume of natural nucleotide polymerase activity assay[42] (PAA) reagents or reversible terminator polymerase activity assay (RT-PAA) reagents were encapsulated by carrier oil (QX200™ Droplet Generation Oil for EvaGreen, Biorad) in monodisperse droplets. Flow rates of 100 µL/h for the two aqueous phases and 800 µL/h for the oil phase were used to generate droplets with a volume of 5 pL at rates of ≈ 10 kHz. The collected droplets were incubated in EP tubes with an overlay of light mineral oil for 5 min at 90 °C to lyse cells, followed by incubation at 55 °C for 2 hours for nucleotide incorporation. For natural nucleotide polymerase activity assay (PAA) reagents preparation, a primer–template complex was annealed in 1×ThermoPol buffer by heating for 5 min at 95 °C and cooling for 5min at 4 °C. The final concentration of DNA primer (PBS2, Table S2), template (ST.1G.-5-Cy3, Table S2) and quencher probe (QP13.-3-Iowa, Table S2) were 1 µM, 0.5 µM and 1.5 µM for annealing, respectively. After annealing, dNTPs (50 µM final) were added. For reversible terminator polymerase activity assay (RT-PAA) reagents preparation, a primer–template complex was annealed in 1×ThermoPol buffer by heating for 10 min at 80 °C and cooling for 1 hours at room temperature. The concentration of primer/template-1 and primer/template-2 (Table S2) were 0.1 µM (final). After annealing, cyanine 5 labeled reversible terminator (0.2 µM final) was added.

**Droplets sorting**

After incubation, the droplets were reinjected into the sorting chip (Figure S1b) at a rate of 10 µL/h. With two streams of spacing oil at a rate of 500 µL/h, the droplets were spaced and enter the detection& sorting area in order. For natural nucleotide polymerase activity selection, a 532 nm excitation laser was focused on the detection& sorting area through a 20× microscope objective (Sunny optical) and the emitted fluorescent light (585/29 nm) was collected using photomultiplier tubes (H10722-20, Hamamatsu), if the fluorescence intensity in a droplet was within the sorting gate (Figure S2a), it was deflected to collection channel and then collected through polyethylene micro-tubing to an Eppendorf tube. For reversible terminator polymerase activity selection, a 532 nm excitation laser was focused on the detection& sorting area, if the ratio of cyanine 5 fluorescence intensity (681/24 nm, collected by photomultiplier tubes through fiber optics) to cyanine 3 fluorescence intensity (585/29 nm, collected by photomultiplier tubes through objective) in a droplet was higher than the set ratio threshold and the cyanine 5 fluorescence intensity (681/24 nm) was higher than the set fluorescence threshold (Figure S2b), it was collected. The positive sorting pulses triggered by MCU-embedded threshold logic triggered the printed circuit board assembly (PCBA) to emit a 20 kHz square-wave signal (50% duty cycle), which was subsequently amplified to 1,000 V via a high-voltage amplifier and transmitted to the field's alloy electrode (Pb-In-Sn eutectic, Tm=75°C) on the sorting chip. This generated dielectrophoretic (DEP) forces, achieving precise droplet deflection into collection channel.

**DNA recovery and amplification**

After droplet sorting, 50uL of preformed droplets made from nuclease free water was added to the collection tube and mixed to ensure the sorted droplets are randomly distributed within the added droplets. Then, the same volume of 1H,1H,2H,2H- perfluorooctanol (Sigma-Aldrich) as the oil in collection tube was added and the tube was vortexed for ≈ 30 s and centrifuged at 2000 r.c.f. for 1 min. The top aqueous layer containing variants gene was transferred into another tube as possible. The oil phase was re-extracted as described above. The demulsification product was used as template for polymerase mutagenic gene amplification with the primers in Table S2 by emulsion PCR to avoid bias. After demulsification, the amplicon was purified by DNA gel purification kit (Tiangen) and assembled with linearized vector by Gibson Assembly Master Mix (New England Biolabs). Product of assembly was purified by DNA Clean & Concentrator-5 (Zymo Research) and transferred into E. coli DH5a electrocompetent cells. The colonies were scraped with a sterile L-shaped spreader after pipetting LB medium onto the overnight plate. Sorted mutagenic plasmid DNA was extracted from the collected cells by plasmid miniprep kit (Tiangen) and transferred into E. coli BL21(DE3) for next round of droplets sorting or microtiter plate screening.

**Lysate activity assay in microtiter plate**

To quantify the activity of reversible terminator polymerase variants, individual E. coli BL21(DE3) colonies were picked and grown in 96-deep-well plates in 300 µL LB medium with 50 µg/mL kanamycin at 37 °C/280 rpm for 12 h. Subsequently, 6 µL of overnight cultures were inoculated to 300 µL of medium and incubated at 37 °C/280 rpm for ≈ 3 h (until an OD600 = 0.6 was reached). Protein expression was induced with IPTG (0.5 mM final; Sangon) and expression for 12 hours at 25 °C and 280 rpm shaking. Cells culture was centrifugated at 3900 g for 20 min, the supernatant was discarded, and cells were washed with 1×ThermoPol buffer and re-suspended with 100 µL lysis buffer (1mg/mL lysozyme in 1×ThermoPol buffer). The cells suspension was transferred to 96-well PCR plates and lysed for 10 min at 37 ℃ and 30 min at 80 ℃ in thermal cycler. Cell lysates was centrifugated at 3900 g for 90 min at 4℃, the supernatant was transferred to a new 96-well PCR plates immediately. For Microtiter plate-based reversible terminator polymerase activity assay, 40 uL of the RT-PAA reagents was added to 10 µL lysate supernatant in 384-well black flat clear bottom microplates. The formation of reversible terminator incorporated primer–template complex was recorded in a plate reader (Tecan) for 2 hours at an excitation wavelength of 530 nm and an emission wavelength of 680 nm. The activity of each variant was determined by the slope of the linear portion of each curve (relative fluorescence units (RFUs) vs. time). Relative activity was calculated by normalizing the sample values to the control.

**Expression and purification of KOD variants and 48°N variants**

Single colony picked from agar plate was grown overnight at 37 °C in volumes of 5 mL LB culture media (containing 50 µg/mL kanamycin). The overnight culture was inoculated in 500 mL LB culture media (containing 50 µg/mL kanamycin) at 37 °C until reaching an OD600 value of 0.6, IPTG was added with a final concentration of 0.5 mM and incubated overnight at 25°C and 280 rpm. The KOD expressed cells were harvested by centrifugation (5000 rpm, 20 min) and resuspended with KOD binding buffer (300 mM NaCl, 20 mM Imidazole, 5% glycerol, 50 mM potassium phosphate, pH 7.4@ 25 °C). Cells were lysed with sonication (400 × 3 s, interval of 5 s) in ice-water bath. Precipitation and supernatant of recombinant cells were separated by centrifugation (16,000 × g, 60 min, 4 °C). The supernatant was applied to a Ni-NTA gravity flow column (2 mL bed volume, Ni Sepharose 6 Fast Flow, Cytiva). The column was washed with 20 column volumes of binding buffer and eluted with KOD elution buffer (300 mM NaCl, 200 mM Imidazole, 5% glycerol, 50 mM potassium phosphate, pH 7.4 @ 25℃). The eluate was concentrated by centrifugation through tubes containing filters with molecular weight cut offs (MWCO) of 30 kDa (Amicon Ultra Centrifugal Filter, 30 kDa MWCO, Merck), after being dialyzed using KOD dialysis buffer (20 mM Tris, 200 mM KCl, 0.2 mM EDTA, 5% glycerol, pH 7.4 at 25 °C) at 4℃ for 18 hours. The purification protocol for 48°N variants was identical to that of KOD DNA polymerase, with the exception of buffer. The composition of the 48°N purification buffer is as follows: 48°N binding buffer (20 mM Tris-HCl, 300 mM NaCl, 20 mM Imidazole, 5% Glycerol, pH 8.5 at 25 °C); 48°N elution buffer (20 mM Tris-HCl, 300 mM NaCl, 500 mM Imidazole, 5% Glycerol, pH 8.5 at 25 °C); 48°N dialysis buffer (40 mM Tris-HCl, 200 mM KCl, 0.2 mM EDTA, 5% Glycerol, pH8.0 at 25 °C). Protein expression was confirmed by SDS–PAGE analysis with coomassie blue staining.

**Assessment of KOD variants on the BGISEQ-500 platform**

We conducted sequencing according to the BGISEQ-500 protocol[63] employing the single-end (SE) mode. For making DNA nanoballs (DNBs), a single-strand circular DNA library (Standard library reagent V3.0, MGI) was PCR-amplified and quantified with Qubit ssDNA Assay Kit (Thermofisher). DNBs (concentration ≥8 ng/µL) was loaded onto the sequencing chip (MGI) with sample preparation machine (BGIDL-50, MGI). For sequencing, the DNA polymerase and substrate for incorporation in standard BGISEQ-500RS high-throughput sequencing set (MGI) were replaced with KOD variants and fluorescently labeled 3'-O-2-[1-(ethyldisulfanyl) ethyl] benzene-1-carbaldehyde modified reversible terminator.

To assess the performance of KOD variants, the purified polymerase was tested throughout a series of 10-cycle sequencing runs at first. The quality metrics used to evaluate the polymerases variants were the "EstErr%" (estimate error rate distribution on reads position), "Lag%" (the ratio of sequence copies to N-1 cycle when sequencing reaches the N cycle, describe the loss of synchrony in the readout of the sequence copies of a DNB, caused by incomplete incorporation of a nucleotide in some portion of DNA strands within DNBs by polymerases), "Runon%" (the proportion of sequence copies that have reached the N+1 cycle), and Q30 (the percentage of reads that pass the Q30 quality filter, i.e. an error rate of less of equal to 0.1%). Those variants that had a better performance in 10-cycle sequencing runs were subjected to single-end 100-cycle sequencing runs.

**Polymerase activity assay with capillary gel electrophoresis**

Primer-template substrates were prepared by annealing a FAM labeled incorporation primer and incorporation template listed in Table S2. When DNA polymerase catalyzes the addition of nucleotide at the 3'-end of primers, reactions were analyzed by capillary gel electrophoresis to identify relevant substrate and product peaks. Fraction of substrate and product was calculated from substrate and product peak areas[58].

$$product\ ratio = \frac{product\ peak\ area}{sum\ of\ substrate\ and\ product\ peak\ areas}$$

**Figure S1:** Design of microfluidic chips for droplet generation and sorting. a) a flow-focusing droplet generation chip (height: 20 μm, colored with grey) with (1) oil phase inlet, (2) water phase inlet for E.coli mixture, (3) water phase inlet for substrate, and (4) outlet for the collection of droplets. b) a dielectrophoresis droplet sorting chip (height: 20 μm colored with grey for matching of droplet diameter and 125μm colored with black for matching of fiber diameter) with (5) inlet for the first spacing oil, (6) inlet for the second spacing oil, (7) inlet for droplet sample, (8) collection for target droplet, (9) waste for non-target droplet, (10) & (11) a pair of electrodes (colored with red and blue) for the control of droplet sorting, one for sorting trigger signal and the other for ground, (12) & (13) a pair of shielding electrodes, (14) channel for installation of fluorescence detection fiber.

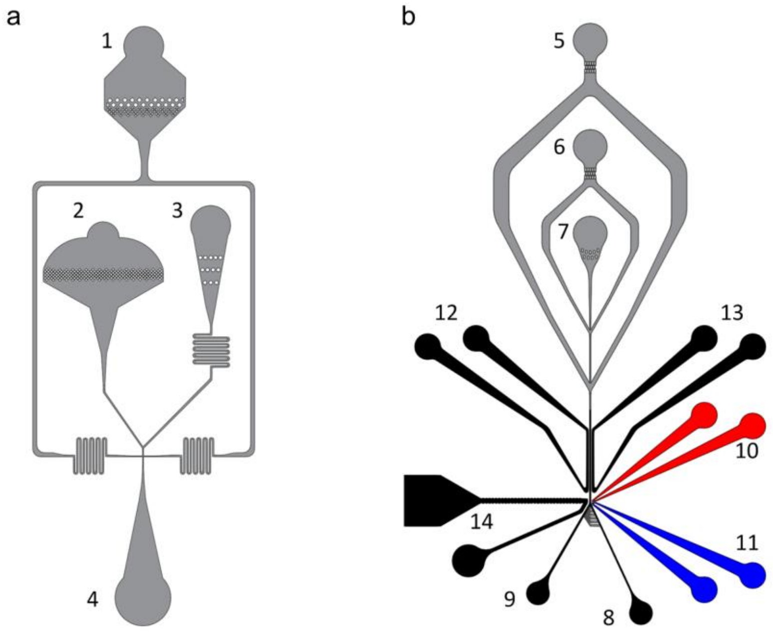


**Figure S2:** To assess the diversity of the constructed library, 20 clones were randomly selected and subjected to sequencing analysis for KOD_M mutant library and 48°N mutant library.

a) KOD_M mutants library clones' sequences.

| clone | L408 | A409 | P410 |
|---|---|---|---|
| 1 | E | E | I |
| 2 | E | N | G |
| 3 | G | A | P |
| 4 | G | G | A |
| 5 | G | H | V |
| 6 | G | V | A |
| 7 | K | D | R |
| 8 | L | V | K |
| 9 | M | G | R |
| 10 | M | H | S |
| 11 | M | L | L |
| 12 | N | S | S |
| 13 | P | E | L |
| 14 | P | L | T |

aKOD_M mutants library clones' sequences. (continued)

| | | | |
|---|---|---|---|
| 15 | R | H | L |
| 16 | R | I | Q |
| 17 | S | S | P |
| 18 | V | L | L |
| 19 | V | Q | P |
| 20 | V | S | G |

b) 48°N mutants library clones' sequences

| clone | L441 | Y442 | P443 | A516 | note |
|---|---|---|---|---|---|
| 1 | | | | | sequencing failure |
| 2 | E | stop coden | G | A | |
| 3 | G | R | S | A | |
| 4 | P | R | R | G | |
| 5 | F | D | H | G | |
| 6 | L | E | S | H | |
| 7 | F | D | W | H | |
| 8 | L | T | K | L | |
| 9 | F | V | M | N | |
| 10 | R | T | G | P | |
| 11 | F | P | S | P | |
| 12 | L | C | M | S | |
| 13 | Q | L | C | T | |
| 14 | R | R | S | T | |
| 15 | L | P | G | V | |
| 16 | R | R | M | V | |
| 17 | stop coden | L | R | V | |
| 18 | V | G | V | V | |
| 19 | A | A | N | Y | |
| 20 | | | | | deletion |

**Figure S3**: The natural nucleotide polymerase activity assay system produces signal when the primer–template complex is extended to a full - length product. The primer–template complex (blue and orange) contains a cyanine3 fluorophore, which is quenched when an Iowa Black FQ - labelled DNA - quencher (black) anneals to the unextended region. This figure is adapted from the study by Larsen et al. published in Nature Communication in 2016 (A. C. Larsen, M. R. Dunn, A. Hatch, S. P. Sau, C. Youngbull, J. C. Chaput, *Nat. Commun.* **2016**, *7*, 1–9.).

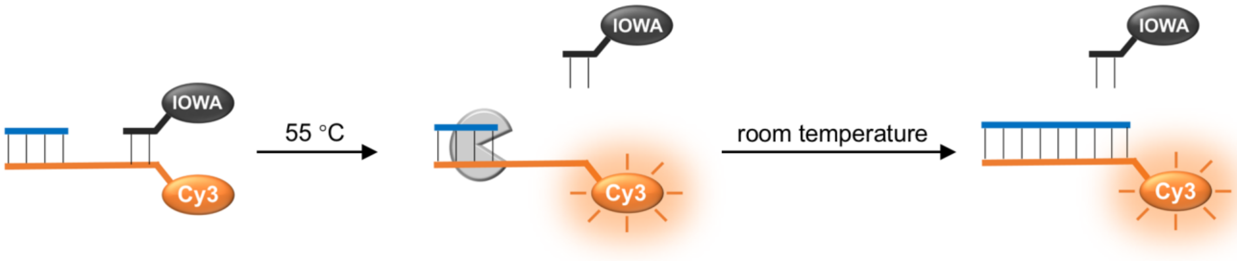


**Figure S4:** Droplets populations plot with the gates applied to select for target subpopulation. Droplets were gated on a) cyanine3 fluorescence intensity/ droplets frequency for natural nucleotide incorporation activity screening and b) cyanine5 fluorescence intensity/ cyanine3 fluorescence intensity for terminator incorporation activity screening.

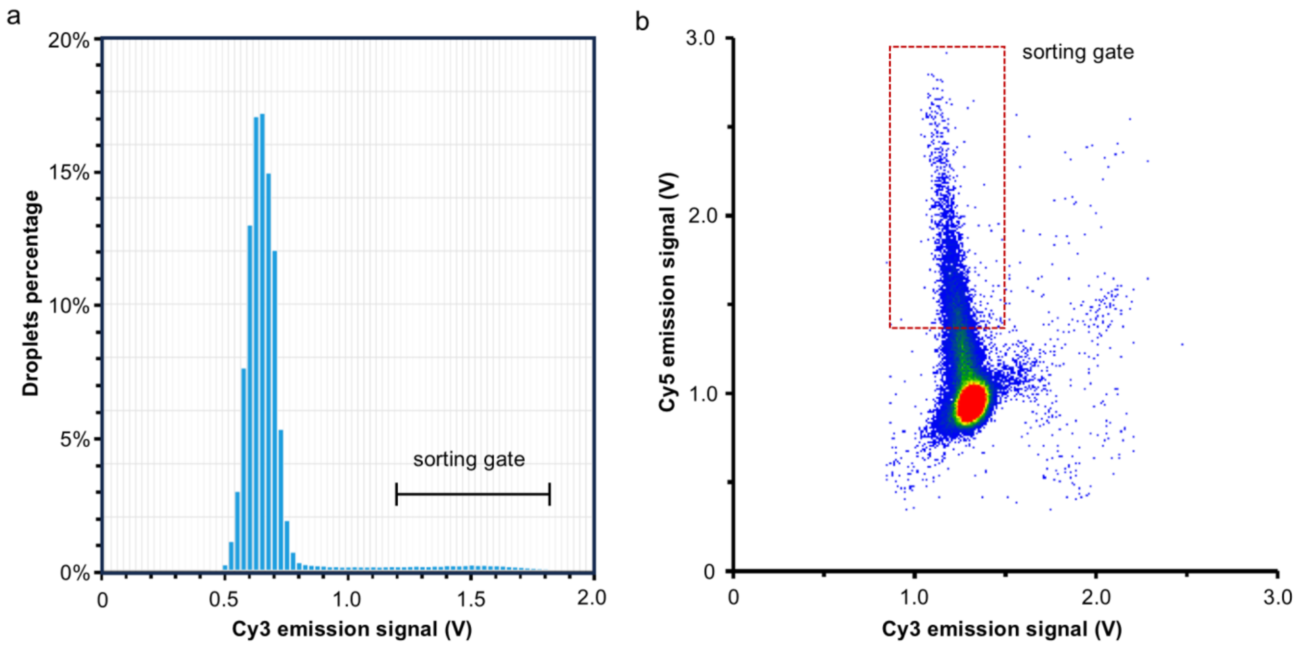


**Figure S5:** The droplets PCR product was separated by 1% TAE-agarose gel electrophoresis and gel-extracted.

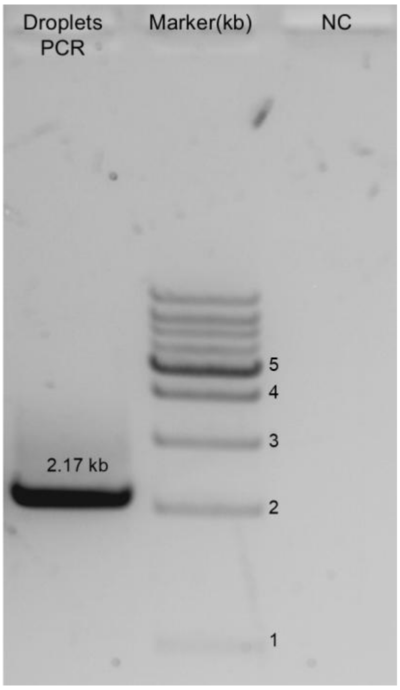


**Figure S6:** Denaturing SDS PAGE for KOD_M and selected variants after Ni-NTA purification and desalting. Lines: 1. KOD_M, 2. KOD_M_L408A/P410I, 3. KOD_M_L408G/P410S, 4. KOD_M_L408G/P410V, 5. KOD_M_L408G/P410A, 6. KOD_M_L408A/A409G/P410G

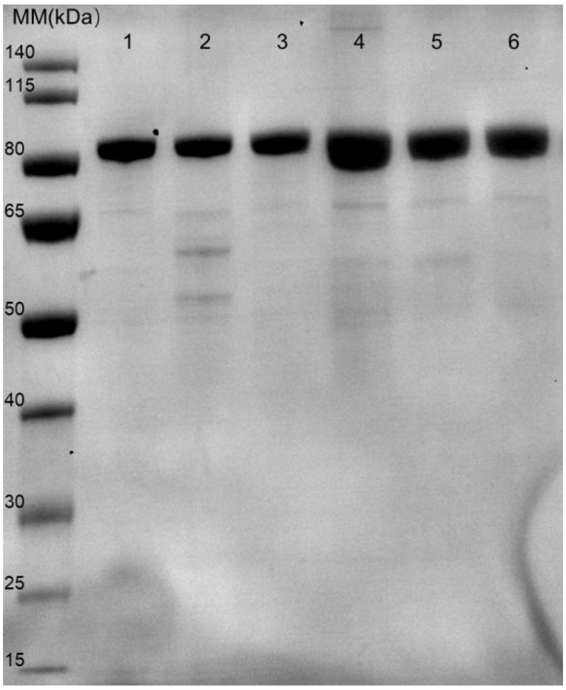


**Figure S7:** Example progress curves of Cy5-labeled SSEB-dTTP polymerization by the KOD_M_L408A/P410I, KOD_M and KOD_C lysate. The increment in relative fluorescence units was recorded at 680 nm via the Tecan Spark™ plate reader.

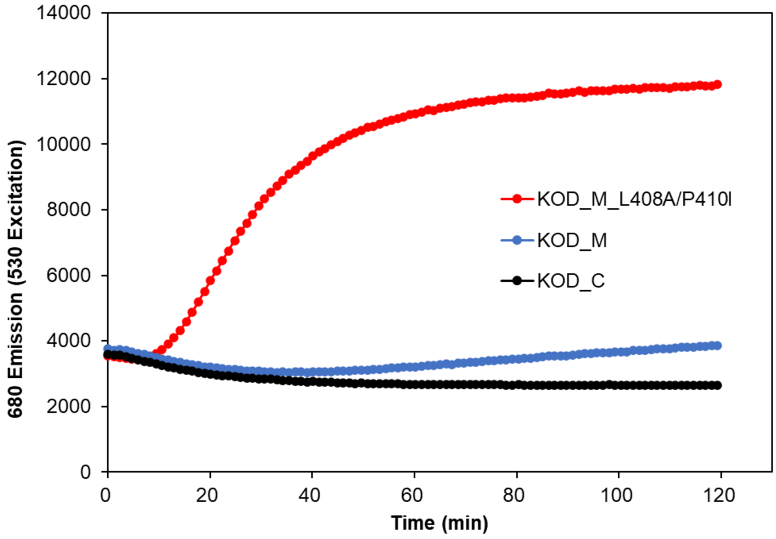


**Figure S8:** Example progress curves of reversible terminator polymerization by the 48°N_D169A/E171A/L441V/Y442A/A516L (48°N_AA_VAL), 27°N_ D153A/E155A/L428V/Y429A/A507L (designated 27°N_AA_VAL), 4°S_ D153A/E155A/L428V/Y429A/A507L (designated 4°S_AA_VAL), 22°S_D147A/E149A/L424V/Y425A/A503L (designated 22°S_AA_VAL) lysate. The increment in relative fluorescence units was recorded at 680 nm via the Tecan Spark™ plate reader.

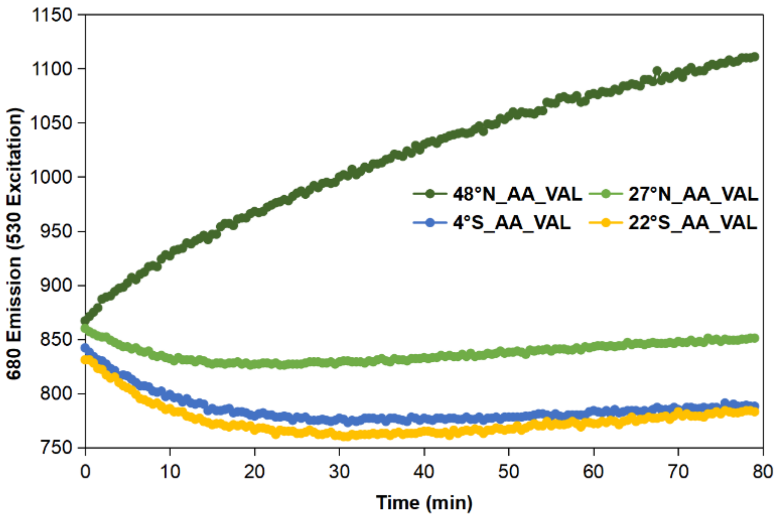


**Figure S9:** Denaturing SDS PAGE for 48°N variants after Ni-NTA purification and desalting. Lines: 1. 48°N_AA_VAL, 2. 48°N_AA_YGLT, 3. 48°N_AA

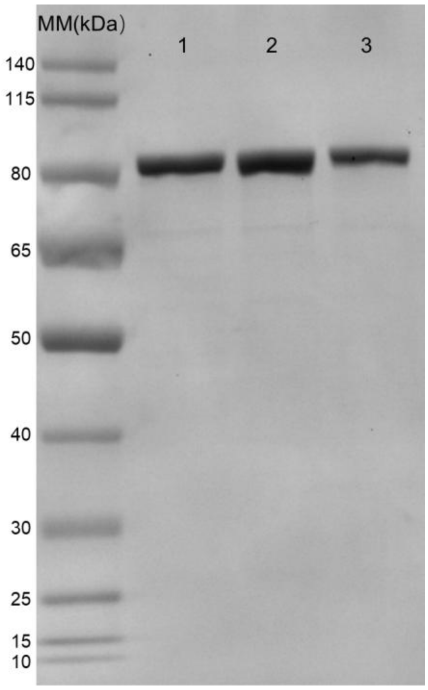


**Figure S10:** 3’-O-azidomethyl-dNTPs incorporation efficiency at 58℃ for 2 min of 48°N_AA_YGLT, revolved by capillary gel electrophoresis

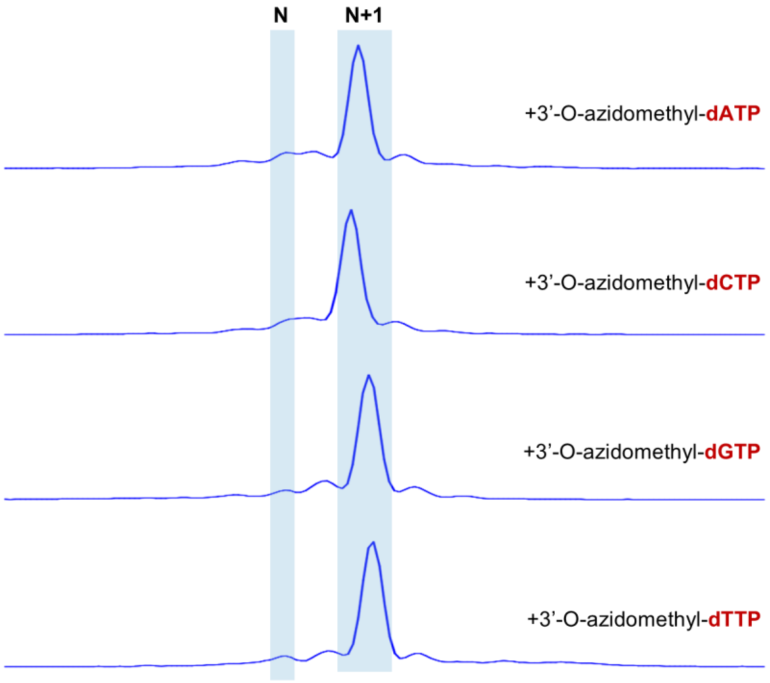


**Figure S11:** Amino acid sequence of KOD DNA polymerase wild-type

ILDTDYITEDGKPVIRIFKKENGEFKIEYDRTFEPYFYALLKDDSAIEEVKKITAERHGTVVTVKRVEKVQKKFLGRP
VEVWKLYFTHPQDVPAIRDKIREHPAVIDIYEYDIPFAKRYLIDKGLVPMEGDEELKMLAFDIETLYHEGEEFAEGPI
LMISYADEEGARVITWKNVDLPYVDVVSTEREMIKRFLRVVKEKDPDVLITYNGDNFDFAYLKKRCEKLGINFALG
RDGSEPKIQRMGDRFAVEVKGRIHFDLYPVIRRTINLPTYTLEAVYEAVFGQPKEKVYAEEITTAWETGENLERVA
RYSMEDAKVTYELGKEFLPMEAQLSRLIGQSLWDVSRSSTGNLVEWFLLRKAYERNELAPNKPDEKELARRRQS
YEGGYVKEPERGLWENIVYLDFRSLYPSIIITHNVSPDTLNREGCKEYDVAPQVGHRFCKDFPGFIPSLLGDLLEE
RQKIKKKMKATIDPIERKLLDYRQRAIKILANSYYGYYGYARARWYCKECAESVTAWGREYITMTIKEIEEKYGFKVI
YSDTDGFFATIPGADAETVKKKAMEFLKYINAKLPGALELEYEGFYKRGFFVTKKKYAVIDEEGKITTRGLEIVRRD
WSEIAKETQARVLEALLKDGDVEKAVRIVKEVTEKLSKYEVPPEKLVIHEQITRDLKDYKATGPHVAVAKRLAARG
VKIRPGTVISYIVLKGSGRIGDRAIPFDEFDPTKHKYDAEYYIENQVLPAVERILRAFGYRKEDLRYQKTRQVGLSA
WLKPKGT*

**Figure S12:** Amino acid sequence of 48°N DNA polymerase wild-type

MTRVVKGFLIDADYETVEGKALIRLYLRDEQGNSFVTYDDSFSPYFYALPGDEPEKVRERILSSDEEGIIQNIAIEEK
RFFGKPVTALRITVHHPQDVPRIRERIRHLEGVDAINEHDILFVRRYLIDRGIKPLTWLELEVEEEQGRLFVRGIEHI
DGELPELKVAAVDIEVYNPKGAPRSKQDPIIMVSMVTSDGMEKVLTWKPVEGEAYVETLSNEKEMLLRFAQLVQE
GDYDVIVGYNTDNFDFPYIKDRVKKLGISLPLGRLNAELEVSRRGALPEARIRGRAHVDLYPIVRRHVKLNSYVLES
VVEELLGIKKEKIDGDKLFTYWDKGGENLAMLARYSLEDARVTLFLAEKFLPLYYELAVIVGQSLNDVARMTSGQL
VEWLLMRFATPRGELIPNHPAGEEYAARASASYVGGYVREPKRGLVENIAVFDFRSLYPSIIVSHNIDPSTLIIGNC
EENRAPGLDYCFSLDKEGFIPEILKGLIQRRAEIKQQMKQVDEDKRRMLSVAEKALKILANSFYGYMGYPRARWY
RRECAESVAAFARMYIKQVMRIAEEDFKLEVVYGDTDSLFVVIPPEKRELAREFLRKVNASMPGMIELEFEGFYKR
GLFVTKKRYALLSEEGKMVVKGLEFVRRDWAPIARETQREVLRILLEEANPEKAAKLVRDVIKRIRERRVTLEDITIY
TQLTKRIKSYKSLEPHVVAAQKLKERGREVAPGMIIGYIITRGTKAISHRALPVEFAKLEDYDPDYYIDNQILPAIQRI
FEAIGYTRDYLREGITQTSLSKWF*

**Figure S13:** Amino acid sequence of 27°N DNA polymerase wild-type

MRGVIIDVDHDISNRDETVIRLFVTDGKQWKMFMEKYRPYFYVVGGELDTVKKALEGAEGVVSVEKVRRKVQWE
EREVLLVRVRHQKYIQRLKEAIEERGAECREHDIPAERKYLYEKGIAPMDVVEVEGDGTWLRSIRKVNGEQELKV
AAIDLEMYARDRMPNPQRDEIIMASYVDSGGKRLVITTKEIDAPFVKKVETEAQLISELNRTVLENDPDIILTYNGDG
FDLPYLKERAHVLGLKIPWGRDGTEPRIRRAGGGSTTVDIVGRAHVDVFRIVQFMAGVGAINTFKLDLENVYKAVL
GKEKVKIEHRDIAKEWREGNLKLLATYNLQDSEACYELGMEFLPLYEELARITASNLYNIVRMSTSQIVEWKLILEA
FREGKLVPKKPKEEEVQRRMMHTYEGAFVKEPVPGLHERIAVLDFRSLYPTIIISHNVDPDTVNAPYCSREDAYVS
PAGHYFCKEPKGLLPKMLEEVLKERFTLKDKMKRMDRKDPEYKIVYAKQQALKIIANSAYGYLGFARARWYSKEC

AEAITAWARKYIRDVMRKAEEFGFRVIYGDTDSVFVVIPEGWGLEKVFEFLEKVNAELPKPMTLEFDGYYVRGIFL
TKRGGKRAAKKKYALIDEKGNLKITGMEYVRRDWAEIAKEVQKRVLELVLGEGKPEEAVNYVKKVIDEIREGKIPK
EKLVIYTQIRRDLDRYEARGPHVAAAMKAMRRGIKIEPGMIIGYIITKRGKSISDKAELAAFVEEGDYDPEYYINNQV
LPAVMPILEELGYTERDIRKGSAQKTLFDFT*

**Figure S14:** Amino acid sequence of 4°S DNA polymerase wild-type

MRGVIIDVDHDISNREETIIRLFLRTETGTTMLQEKYRPYFYVVGEELDAAEAELKKAEGVESVERVKRYVRWEEK
ELLLVRVHHQKYIPKIKERLEELNVECREHDIPVERKYLYEKGIVPFGVVEFTEKNGWITEIKPAHGEVPIRVAAFDL
EMYARDRMPDPKKDEIIMASYVDSEGNKIVITTKKIDAPFVKTVETEEQLISELTRLIKENDPDIILTYNGDAFDLPYL
KERAHVLGLRIPWGRDGSEPRIRKAGGGNTTVEITGRAHVDVFQIVQFMAAVGAINTFKLDLENVYKTVLGREKV
KIEHREIANVWAKGDLNELALYNLQDSEACYELGMEFLPLYAELARITASNLYNVVRMSTSQIVEWKLILEAFRAKK
VVPRKPREEEVRTRLTQSYEGAFVREPIPGLHENIVVLDFRSLYPTIIISHNVDPDTVNAPYCTREEAYVSPAGHYF
CKEPKGLLPRTLEDVLGERFRLKDQMKKMDKKNPQYRIVYAKQQALKIIANSAYGYLGFARARWYSRECAEAITA
WARKYIKDVMKWAEDKGFTVLYGDTDSVFLVLPKGWGVDKALEFMEEVNKRLPEPMSLEFDGYYVRGIFLTKRG
GKKAAKKKYALIDERGNLKITGMEYVRRDWAEIAKDVQRKVLELVLGEGKPEEAVKYVRKVIDKIRAGKVPKEKLII
YTQIRRDLDRYEARGPHVAAAEKAMQRGIKIEPGTIIGYIVTKRGKSISDKAELAAFVEDGDYDAEYYINNQVLPAV
MPILEELGYTERDISIGSAQKTLLDFT*

**Figure S15:** Amino acid sequence of 22°S DNA polymerase wild-type

MRGVIVDVDHDLGNREQTIIRLYVRGEDGSVHVIRKEYTPYFYVLSENPEDTVQKIRELSGVKSAEIVERFVRWEK
KRVILVRVAHQKYINRIREKIKEMGGECREHDINVERKFLYEHGLEPLGGIDTETLSPVETRVEPVIAAFDLEVYAKD
RMPEPDRPENEIIMASWVDSTGRRVVITTKDIERDYVVKVETEAQIISELNRLMREVDPDIVLTYNGDNFDLPYLRE
RARFLGMRIPWGRDGSEPRIRRGPTGNTTVDIAGRAHVDVFQIVQFMAGIGAINSIHLDLESVYEAVLGKKKVKIK
HMDIADQWRSGDLGLLADYNLQDSIACYELGMEFLPLYVELSRITVSPLPNTTRMSASQLVEWTLIRRAFKMGKV
VPRKPREEEVRARMMETYQGAFVKEPVPGLHERIVVLDFRSLYPTIIISHNVDPDTANAPYCSRENAYVSPVGHY
FCKEPKGLIPGMLEEVLELRFALKDKLKKMDKDDPDYKPLYAKQKALKIVANATYGYLGFARARWYCKVCAEAVT
AWARHYIKKVMKWAEEHGFTVIYGDTDSVFIKLGEDQTLEDALAFMGWVNERLPKPMALDFDGYYVRGIFLSKR
RSGKAAKKKYALIDEKGRLKITGLEYVRRDWAEIAKEVQAKVLEYVLKEGDVKKAVDYVKRVIADVRAGKIPKEKLII
YTQIQKELDRYAARGPHVAAALKAMKRGVHIEPGMIIGYIITKRGSSISDKAELAQFVREGDYDPEYYVEHQIIPAVL
PILEELGYDERALKQDAGQRSIFDFT*

**Table S1. Strains and plasmids used in this study.**

| Strains and plasmids | | |
|---|---|---|
| E. coli DH5a | Host for cloning plasmids | Lab stock |
| E. coli BL21 (DE3) | Host for expression plasmids | Lab stock |
| E. coli BL21-KOD | E. coli BL21 (DE3) carrying plasmid pD441-KOD | Lab stock |
| E. coli BL21-48°N | E. coli BL21 (DE3) carrying plasmid pET28a-48°N | Lab stock |
| pD441-KOD | pD441 harboring the KOD gene | Lab stock |
| pET28a-48°N | pET28a harboring the 48°N gene | Lab stock |

**Table S2. Oligonucleotides used in this study.**

| oligonucleotides | Sequence of oligonucleotides |
|---|---|
| For generation of KOD mutant | |
| D141A/E143A-F | 5'-TGCTGGCGTTCGCCATCGCCACTCTGTACCACG-3' |
| D141A/E143A-R | 5'-CGTGGTACAGAGTGGCGATGGCGAACGCCAGCA-3' |
| L409A-F | 5'-ATTTGGATTTTCGTAGCTTGGCCCCGAGCATCATTATCACGC-3' |
| L409A-R | 5'-GCGTGATAATGATGCTCGGGGCCAAGCTACGAAAATCCAAAT-3' |
| S451T-F | 5'-CGGGCTTTATCCCGACCCTGCTGGGTG-3' |
| S451T-R | 5'-CACCCAGCAGGGTCGGGATAAAGCCCG-3' |
| A485L-F | 5'-GACTACCGTCAACGTCTGATCAAGATCCTGGCGAA-3' |
| A485L-R | 5'-TTCGCCAGGATCTTGATCAGACGTTGACGGTAGTC-3' |

**Table S2. Oligonucleotides used in this study.(continued)**

| | |
|---|---|
| For generation of KOD combinatorial mutagenesis libraries | |
| L408/Y409/P410-gene-F | 5'-GTCTATTTGGATTTTCGTAGCVNKVNKVNKAGCATCATTATCACGCATAA-3' |
| L408/Y409/P410-gene-R | 5'-ATACGAATTCGCCAGGATCTTGATCAGACGTTGACGGTAGTCCAGTAATT-3' |
| L408/Y409/P410-vector-F | 5'-GCTACGAAAATCCAAATAGACGATGTTCTCC-3' |
| L408/Y409/P410-vector-R | 5'-AAGATCCTGGCGAATTCGTATTATGGT-3' |
| For amplifier of KOD mutants gene from droplets demulsification product | |
| KOD-gene-F | 5'-TACTTCTATGCTCTGCTGAAAGACGATAGCGCGATTGAGG-3' |
| KOD-gene-R | 5'-TTTTCTGGTAACGCAGATCTTCCTTGCGGTAACCGAAGGC-3' |
| For amplifier of linerized pD441 vector | |
| pD441-vector-F | 5'-TTTTCTGGTAACGCAGATCTTCCTTGCGGTAACCGAAGGC-3' |
| pD441-vector-R | 5'-CCTCAATCGCGCTATCGTCTTTCAGCAGAGCATAGAAGTA-3' |
| For generation of 48°N mutant | |
| D169A/E171A-F | 5'-GAAAGTGGCGGCGGTTGCTATTGCAGTGTATAACCCGAAAG-3' |
| D169A/E171A-R | 5'-CTTTCGGGTTATACACTGCAATAGCAACCGCCGCCACTTTC-3' |
| For generation of 48°N combinatorial mutagenesis libraries | |
| L441/Y442/P443/A516-gene-F | 5'-GTGTTTGATTTTCGCAGCNNKNNKNNKAGCATTATTGTGAGCCATAAC-3' |
| L441/Y442/P443/A516-gene-R | 5'-CTGTTCGCCAGAATTTTTAAMNNTTTTTCCGCCACGCTCAGC-3' |
| L441/Y442/P443/A516-vector-F | 5'-TTAAAAATTCTGGCGAACAGCTTTTATGGCTATATG-3' |
| L441/Y442/P443/A516-vector-R | 5'-GCTGCGAAAATCAAACACCGCAATATTTTCCAC-3' |
| For amplifier of 48°N mutants gene from droplets demulsification product | |
| 48°N-gene-F | 5'-TACGACTCACTATAGGGGAATTGTGAGCG |
| 48°N-gene-R | 5'-TGTTAGCAGCCGGATCTCAGTGG |
| For amplifier of linerized pET28a vector | |
| pET28a-vector-F | 5'-AGATCCGGCTGCTAACAAAGCC-3' |
| pET29a-vector-R | 5'-TCCCCTATAGTGAGTCGTATTAATTTCGCG-3' |
| For natural nucleotide polymerase activity assay | |
| ST.1G.-5-Cy3 | 5'-Cyanine3-ACAACCATACTCTCCTCATCACTATTCAACTTACAATCGATACAACCTTATAATCCACATGGCTACTGCATACGAGTGTC-3' |
| QP13.-3-Iowa | 5'-AGAGTATGGTTGT-IABkFQ-3' |
| PBS2 | 5'-GACACTCGTATGCAGTAGCC-3' |
| For reversible terminator polymerase activity assay in microtiter plate | |
| primer/template-1 | 5'-Cyanine3-CGTGTAAGCGTAATAGGATCCCGACTCACTATGGACG-3' |
| primer/template-2 | 5'-Cyanine3-Cy3-CGTGTAACGTCCATAGTGAGTCGGGATCCTATTACGC-3' |
| For reversible terminator polymerase activity assay with capillary gel electrophoresis | |
| FAM-Incorporation primer | 5'-FAM-AGTGAATTCGAGCTCGGTACCCGGGGATCCTCTAGAGTCGACCTGCAGGC-3' |
| Incorporation template(+A) | 5'-TTGCTCGTTTGCTGGGTGCCTGCAGGTCGACTCTAGAGGATCCCCGGGTACCGAGCTCGAATTCACT-3' |
| Incorporation template(+C) | 5'-TTGCTCGTTTGCTGGGGGCCTGCAGGTCGACTCTAGAGGATCCCCGGGTACCGAGCTCGAATTCACT-3' |
| Incorporation template(+G) | 5'-TTGCTCGTTTGCTGGGCGCCTGCAGGTCGACTCTAGAGGATCCCCGGGTACCGAGCTCGAATTCACT-3' |

**Table S2. Oligonucleotides used in this study.(continued)**

| | |
|---|---|
| Incorporation template(+T) | 5'-TTGCTCGTTTGCTGGGAGCCTGCAGGTCGACTCTAGAGGATCCCCGGGTACCGAGCTCGAATTCACT-3' |
